# Transcriptomics uncovers outer membrane remodeling as conserved strategy for *Burkholderia cenocepacia* survival in protozoan and macrophage phagosomes

**DOI:** 10.64898/2026.10.08.756501

**Authors:** Álvaro Morón, Alaa E. Tarhouchi, Iván Belinchón, Nerea Castañeda, Patricia de Francisco, Silvia Díaz, Francisco Amaro

## Abstract

Bacterivorous protists expose bacteria to hostile stress conditions that resemble those encountered within macrophage phagosomes and may therefore act as training grounds for bacterial pathogens. However, the mechanisms enabling bacterial survival within protozoan phagosomes remain poorly understood. Here we characterized the transcriptome of the opportunistic pathogen *Burkholderia cenocepacia* during transit through phagosomes and expelled food vacuoles of the ubiquitous protozoan ciliate *Tetrahymena elliotti*. Our data revealed a coordinated response involving bacterial metabolic reprogramming, with the upregulation of NADPH-generating pathways and uptake systems for host-derived nutrients, alongside the activation of several stress responses and envelope remodeling to maintain outer membrane barrier function and repair phagosomal damage. Disruption of antioxidant defenses, the RpoE-dependent envelope stress response, the Mla pathway and other proteins associated with outer membrane integrity severely impaired intracellular survival in both protozoa and macrophages, revealing conserved mechanisms required for persistence within evolutionarily distant phagocytes. This adaptive response not only contributed to bacterial survival but it also primed *B. cenocepacia* for increased chlorhexidine resistance and adhesion to collagen and human lung epithelial cells. Collectively, our findings reveal that protozoan phagosomes drive physiological adaptations with consequences extending beyond grazing resistance. Protozoa may therefore help disseminate stress-adapted bacteria with enhanced resistance and colonization potential, providing a mechanistic link between adaptation to environmental predators and survival in the human host.

## Introduction

Protozoan predation is now recognized as a major selective pressure shaping the evolution of opportunistic bacterial pathogens [1]. This hypothesis stems from the fact that protozoan grazing represents a primary cause of bacterial mortality across natural and anthropogenic environments [2], and that some bacterial defenses originally evolved against protist predators (e.g., mechanisms to resist lysosomal digestion or production of anti-eukaryotic toxins) may act as virulence factors during infection in humans [3–5]. Furthermore, bacteria that evade intracellular digestion can exploit their protist predators as protective reservoirs in which they may multiply, or as training grounds where to increase their resistance towards stress and antimicrobials [6–8]. Bacterivorous protists (protozoa such as amoebae or ciliates) and macrophages use the same conserved mechanisms to ingest and kill bacteria. Protozoa capture bacteria into food vacuoles (phagosomes) that are acidified and exposed to oxidative radicals (reactive oxygen species or ROS, and reactive nitrogen species or RNS), osmotic stress, antimicrobial peptides and digestive enzymes [4]. Bacteria that survive digestion are eventually released as free cells or packaged into membrane vesicles named expelled food vacuoles (EFVs), which range from 1 to 5 μm in diameter and could thus serve as respirable vehicles for airborne pathogen dissemination [9]. Indeed, it has been shown that passage through protozoa and subsequent packaging into EFVs enhances bacterial resistance to stress and antibiotics, and even virulence [7,8,10].

*Burkholderia cenocepacia* is a Gram-negative that belongs to the *Burkholderia cepacia* complex (Bcc), a group of 28 closely related species ubiquitous in both natural and anthropogenic environments that can also cause life-threating infections in immunocompromised people [11,12]. Bcc infections are particularly problematic in cystic fibrosis (CF) patients, where they are associated with accelerated decline in lung function, as well as higher morbidity and mortality compared to other infectious agents [13]. Over the past decades, Bcc bacteria have also been linked to hospital-associated outbreaks, most often caused by contaminated medical products such as chlorhexidine solutions [14,15]. We have recently shown that *B. cenocepacia* survives grazing by ciliates found in the natural environment and water distribution networks, and that passage through protozoan phagosomes enhances bacterial resistance to stress and antimicrobials [7]. While this phenomenon has been observed for a variety of pathogenic bacteria [8–10], the underlying molecular mechanisms remain poorly characterized. This represents a critical knowledge gap, as packaging in EFVs appears to play an important role in the persistence, dissemination and infectious potential of bacterial pathogens. Furthermore, elucidating the mechanisms that enable bacteria to survive phagocytosis by protists may reveal conserved strategies for intracellular persistence. Here, we analyzed the *B. cenocepacia* transcriptome during its transit through phagosomes and EFVs of the ubiquitous ciliate *Tetrahymena elliotti.* We show that *B. cenocepacia* activates a coordinated network of stress responses and an outer membrane (OM) remodeling program that facilitate intracellular survival in protists and macrophages while enhancing bacterial antimicrobial resistance and adherence to human epithelial cells. Overall, this study identifies mechanisms responsible induced adaptive tolerance by means of passage through protozoan phagosomes that may favor pathogen dissemination and survival in the human host.

## Materials and Methods

### Microorganisms, cell lines and culture conditions

*B. cenocepacia* and *E. coli* were grown in lysogeny broth (LB) medium at 37 °C. The bacterial strains used in this study are listed in Table S1. When required, antibiotics were used at the final concentration: 50 μg/ml gentamicin (Panreac) for *B. cenocepacia,* 100 μg/ml trimethoprim (Panreac) for *B. cenocepacia* and 50 μg/ml for *E. coli*, 150 μg/ml tetracycline (Panreac) for *B. cenocepacia* and 20 μg/ml for *E. coli*, and 40 μg/ml kanamycin (Panreac) for *E. coli*.

*T. elliotti* strain 4EA was grown at 28 °C in 2% proteose peptone (Pronadisa) supplemented with 10 μM FeCl_3_ (Sigma Aldrich) and 250 μg/ml of both streptomycin sulfate and penicillin G (Sigma Aldrich). THP-1 monocytes (ATCC number TIB-202) were cultured at 37 °C with 5% CO_2_ in RPMI-1640 medium (Gibco) supplemented with 10% heat-inactivated fetal bovine serum (FBS, Gibco) and 100 μg/ml of both streptomycin and penicillin (Sigma Aldrich). THP-1 monocytes were differentiated to adherent macrophages by supplementing media with 75 ng/ml phorbol 12-myristate 13-acetate (PMA) (Sigma Aldrich) at 37 °C with 5% CO_2_ for 2 days before infection.

Human lung epithelial A549 cells (ATCC number CCL-185) were cultured at 37 °C with 5% CO_2_ in Dulbecco’s Modified Eagle (DMEM) medium (Biowest) supplemented with 10% FBS and 100 μg/ml of both streptomycin and penicillin.

### Construction of mutant and reporter strains

Unmarked non-polar deletion mutants were constructed by using the allelic replacement approach developed by Flannagan and co-workers [16]. Upstream and downstream flanking regions of each target gene were PCR amplified from *B. cenocepacia* K56-2 gDNA with Q5 High-Fidelity DNA polymerase (New England Biolabs) and primers listed in Table S2. Plasmids were introduced into *B. cenocepacia* by triparental mating from an *E. coli* donor strain and using the helper strain *E. coli* HB101 carrying plasmid pRK2013 as we described in [7]. Deletion of target genes in each generated mutant strain was confirmed by PCR using primers listed in Table S3.

### Bacterial RNA isolation and RNA sequencing

Co-cultures protist:bacteria (1:100) were established in 0.01 M Tris-HCl pH 7.5 buffer and incubated at 28 °C. After 2, 4 or 24 h, bacteria were released from *T. elliotti* food vacuoles by lysing ciliates on ice for 20 min in an RNA stabilization solution: 1% acidic phenol and 19% ethanol in DEPC-H_2_O supplemented with 0.1% SDS (all from Fisher Scientific). Bacteria were then collected by centrifugation (3500 x g, 10 min) and total RNA was extracted and DNAseI-treated using the Direct-zol RNA miniprep plus kit (Zymo Research). As control, total RNA was isolated from bacteria incubated for 2, 4 and 24 h in 0.01 M Tris-HCl pH 7.5 in the absence of ciliates by following the same procedure. Three biological replicates were analyzed for each condition at 2 h, and two biological replicates for each condition at 4 and 24 h. RNA quality was evaluated by measuring the rRNA integrity through capillary electrophoresis in the Agilent 2200 Tape Station Systems (Agilent Technologies). Samples with an RNA Integrity Number (RIN) ≥ 7.5 were selected for further analysis. Libraries were prepared with the Illumina Stranded Total RNA Prep with Ribo-Zero Plus (Illumina Technologies) by following manufacturer’s directions. The quality of libraries was validated by Qubit dsDNA HS Assay Kit (Thermo Fisher) and Agilent 2200 Tape Station System. Libraries from RNA samples isolated at 2 h were sequenced on an Illumina NovaSeq 6000 at the Translational Genomics NGS and Bioinformatics Service at the Ramón y Cajal Health Research Institute (IRYCIS), producing 144.41 million 70-bp single-end reads (Table S4). Libraries from RNA samples isolated at 4 and 24 h were sequenced on an Illumina NextSeq Mid Output, at the López-Neyra Institute of Parasitology and Biomedicine (IPBLN-CSIC), producing 128.49 million 75-bp paired-end reads (> 11.6 million reads per sample) (Table S4).

### Analysis of differential gene expression

Reads were quality assessed with FastQC [17] and rRNA reads were filtered out with SortmeRNA v4.3.6 [18]. Reads were processed and mapped to the *B. cenocepacia* K56-2 genome sequence (GeneBank accession number CP053300-3) [19] using Burrows-Wheeler Aligner (BWA) [20], yielding 231 million genome-mapped reads for further analysis. The number of read counts for feature was obtained using featureCounts v.2.12.3 [21]. Differentially expressed genes (DEG) were calculated using the Bioconductor package DESeq2 v1.38.0 [22] in R. A gene was considered differentially expressed if its false discovery rate (FDR)-corrected p-value were below 0.05 and the absolute log2-fold change (log2FC) was greater than 1 (corresponding to a minimum of 2-fold upregulation or downregulation). Principal component analysis (PCA) was carried out with the plotPCA function of DESeq2 to test whether biological replicates cluster and whether there is segregation according to treatments. DEG were further functionally categorized using the Cluster of Orthologous Group (COG) database [23]. COG functional categories were assigned to *B. cenocepacia* K56-2 genes based on COG annotations of their homologs in *B. cenocepacia* J2315 retrieved from the Burkholderia Genome Database [24], and annotations generated with eggNOG-mapper v2 [25]. COG category enrichment analysis among up- and downregulated genes was assessed using a hypergeometric test implemented with the phyper function in R, with all protein-coding genes as the reference background. Gene Ontology (GO) enrichment analysis using the GO database [26] was carried out using the R package topGO v2.50 [27] with the Fisher’s exact test and the weight01 algorithm. We focused on the biological process category of GO terms. Kyoto Encyclopedia of Genes and Genomes (KEGG) [28] terms were assigned using the web server DAVID [29]. Venn diagrams and heatmaps of DEG were built with R packages VennDiagram v1.7.3 [30] and ComplexHeatmap v2.14 [31], respectively.

### RT-qPCR

RNA was isolated from synchronized log-phase (OD_600_ = 0.3) bacterial cultures growing in LB medium with the Direct-zol RNA miniprep plus kit (Zymo Research) as we described in [7]. Total RNA was DNAseI-treated (Zymo Research) and retrotranscribed with the Maxima Reverse Transcriptase (Thermo Scientific) by following manufacturer’s directions. The resulting cDNA samples were amplified with specific primer pairs in 96-well microtiter plates 4titude® using 1x TB Green Premix Ex Taq (Takara) as previously described [7]. The mRNA expression levels were quantified using the 2^-ΔΔCt^ method and normalized to the housekeeping gene *gyrA* (locus tag K562_RS14125). Both non-template controls (NTC) and RT-minus controls showed no amplification.

### *Burkholderia* intracellular survival in ciliates and macrophages

Intracellular survival in *T. elliotti* was performed as we described in [7]. Briefly, synchronized bacterial cultures in log-phase (OD_600_=0.3) were washed and mixed with ciliates (3x10^5^ cells/mL) in a 1:100 (protist:bacteria) ratio in 0.01 M Tris-HCl pH 7.5 buffer and maintained at 28 °C. At times 0 and 24 h protists were lysed by adding 1% Triton X-100 (Sigma Aldrich). The number of viable bacteria were quantified by CFU counts on LB agar plates. For each strain, at least three independent experiments were performed.

Bacterial survival and replication in THP-1 macrophages was determined as follows. Synchronized (OD_600_ = 0.3) bacterial cultures washed and resuspended in RPMI-1640 medium were used to infect THP-1 macrophage monolayers at a multiplicity of infection (MOI) of 2, in 24-well plates. Plates were centrifuged for 5 min at 400 x g to synchronize infection (time 0) and then incubated at 37 °C in 5% CO_2_ for 2 h to allow bacterial entry. Then, at 2 h post infection (2 hpi), infected macrophages were washed 3 times with Dulbecco’s Phosphate Buffered Saline (DPBS, Biowest) and treated for 2 h with RPMI-1640 supplemented with 1 mg/mL of both amikacin and ceftazidime (Sigma Aldrich) to kill any extracellular bacteria. To quantify the number of internalized bacteria, at 4 hpi macrophages were washed 3 times with DPBS and lysed with 0.5% Triton X-100 in DPBS. Lysates were serially diluted in PBS and plated onto LB agar plates for CFU enumeration. To quantify bacterial intracellular growth, at 4 hpi macrophages were washed 3 times with DPBS and incubated in fresh RPMI-1640 supplemented with 10% FBS and 100 μg/ml of gentamycin for up to 20 h at 37 °C with 5% CO_2_. At 20 hpi, macrophages were washed with DPBS and treated for 2 h with amikacin and ceftazidime (1 mg/mL each) to kill extracellular bacteria. Then, macrophages were again washed 3 times with DPBS, lysed with 0.5% Triton X-100 and serial dilutions were plated on LB agar plates for CFU enumeration. An aliquot of the third wash with DPBS was plated to assess the efficacy of the antibiotic treatment. Greater than 99.9% of bacteria were killed by the antibiotic treatment. The relative fold replication of intracellular bacteria was calculated as “replication index” as the CFU recovered at 20 hpi relative to those recovered at 4 hpi.

### Bacterial adhesion assays

Co-cultures protist:bacteria (1:100 ratio) were established as outlined above. After 24 h at 28 °C, EFVs were purified as previously described [7]. Bacteria were then released by adding 1% Triton-X100 and washed twice with 0.01 M Tris-HCl pH 7.5 buffer. Bacteria incubated in the same conditions but in the absence of ciliates were identically processed in parallel and served as controls. Bacteria were then washed three times, resuspended in DMEM medium and added to A549 monolayers (5 x 10^5^ cells per well in a 24-well plate) at MOI of 50, centrifuged for 5 min a 400 x g and incubated at 37 °C for 1 h Afterwards, non-adherent bacteria were removed by washing each well three times with DPBS. A549 cells were then lysed by adding 0.5% Triton-X100 and serial dilutions were plated on LB agar plates to determine the number of adhered bacteria by CFU counting. The percentage of adherent bacteria was calculated as the number of adherent bacteria relatively to the initial bacterial load added to the A549 monolayer.

Bacterial adherence to Millicoat® collagen-I-coated 96-well plates (Sigma Aldrich) were assessed by following a similar protocol. Bacteria released from EFVs or control bacteria were washed three times, resuspended in PBS and added to collagen-coated plates (5 x 10^6^ bacteria/well). Plates were centrifuged at 400 x g for 5 min and incubated at 37 °C for 1 h. Afterwards, wells washed three times with PBS and treated with 0.5% Triton X-100 in PBS for 1 h to desorb the bound bacteria. Serial dilutions were plated on LB agar plates to determine the number of adhered bacteria.

### Membrane permeability assay

Changes in the OM permeability were assessed with the N-phenyl-1-naphthylamine (NPN), a lipophilic dye that is weakly fluorescent in aqueous environments but becomes highly fluorescent in hydrophobic environments such as the inner leaflet of the OM [32]. Bacteria in late log phase (OD_600_=1) were washed twice and incubated for 30 min at room temperature in 5 mM HEPES buffer pH 7.2 with 10 mM NaN_3_ (Sigma Aldrich). An equal volume of cells was mixed with HEPES buffer supplemented with 80 μM NPN (Across Organics) and transferred to 96-well black plates. The fluorescence signal was measured using the Tecan Infinite® 200 PRO plate reader with filter sets Excitation 360/40 nm and Emission 420/40 nm. 30 μg/ml chlorhexidine gluconate (Thermo Scientific Chemicals) was used as positive control for membrane permeabilization. NPN fluorescence in HEPES buffer alone and bacterial fluorescence in the absence of NPN were measured to account for background fluorescence and cell autofluorescence, respectively. For each strain, the NPN uptake ratio was calculated by dividing net NPN-associated fluorescence in the bacterial suspension (fluorescence of NPN-stained bacteria minus cell autofluorescence) by the corresponding net NPN fluorescence in HEPES buffer (fluorescence of NPN in buffer minus buffer alone).

### Assessment of chlorhexidine resistance of bacteria recovered from EFVs

Bacteria released from EFVs and control bacteria incubated in the absence of protists were obtained as described above. Both populations were exposed to 10-30 μg/ml chlorhexidine gluconate in 0.9% NaCl solution for 24 h at 37 °C. Afterwards, bacteria were serially diluted in fresh 0.9% NaCl and plated on LB agar plates. The number of surviving bacteria was estimated by CFU counts. Three independent experiments were performed.

### Statistical analysis

Statistical analyses were performed with GraphPad Prism v.8 (GraphPad software). The statistical tests applied to each dataset, together with the corresponding significance thresholds, are indicated in the figure legends. Statistical methods applied to transcriptomic and other bioinformatic datasets are described in the respective sections.

### Data availability

Raw sequencing files can be found under accession number BioProject PRJNA1497282 at NCBI https://www.ncbi.nlm.nih.gov/.

## Results

### Transit through protozoan phagosomes rewires the *B. cenocepacia* transcriptome

To gain deeper insight into the conditions encountered by *B. cenocepacia* within *T. elliotti* phagosomes and the mechanisms enabling bacterial survival and enhanced stress resistance following transit through protists, we analyzed the transcriptome of *B. cenocepacia* K56-2 recovered from protozoan food vacuoles at 2, 4, and 24 h post-feeding (Fig. 1 A). The 2 h time point was selected as it corresponds to the time required for respiratory burst and acidification in *Tetrahymena* food vacuoles [33]. Time points 4 and 24 h capture bacterial transcriptome within late-stage and expelled food vacuoles (EFVs), respectively. The principal component analysis (PCA) revealed that RNA samples extracted from bacteria recovered from protozoan food vacuoles clustered separately from the control RNA samples extracted from bacteria incubated in the absence of ciliates, indicating differences in their expression profile and underscoring the impact of protozoan phagosomes on the bacterial transcriptome profile (Fig. 1 B and S1). Compared to planktonic bacteria maintained in the absence of ciliates, 4468 genes (64.1 % of K56-2 protein-coding sequences) were differentially expressed at 2 h (FDR-adjusted p < 0.05) (Fig 1 C-D). Among these, 1896 (27.2 %) were upregulated and 2020 (29 %) were downregulated by at least two-fold (log2FC ≥ |1| and FDR-adjusted p-value < 0.05) (Fig 1 C-D). At 4 h, 2286 genes (32.8 %) showed altered expression, with 999 (14.3 %) upregulated and 1086 (15.3 %) downregulated (Fig. 1 C-D). By 24 h, 1933 genes (27.7 %) were differentially expressed, with 821 (11.8 %) upregulated and 934 (13.4 %) downregulated (Fig. 1 C-D). Overall, these results indicate that passage through protozoan phagosomes triggers extensive and dynamic transcriptional changes in *B. cenocepacia*. A complete list of differentially expressed genes (DEG) during passage through *T. elliotti* (at 2, 4 and 24 h) is provided as supporting information (Table S5).

**Fig. 1.**
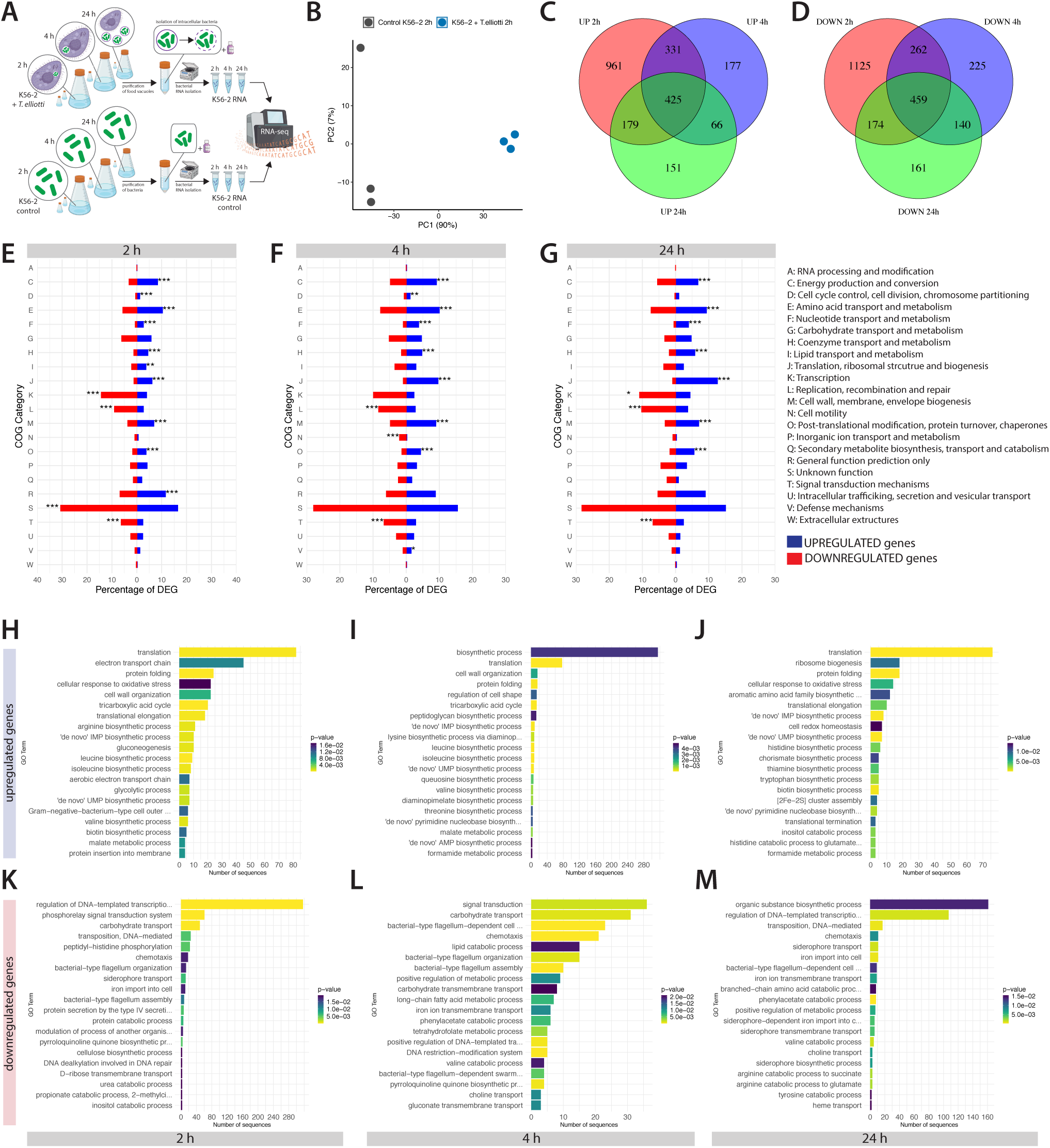
Global differential gene expression analysis of *B. cenocepacia* K56-2 during transit through *T. elliotti*. **(A)** Experimental workflow. RNA-seq analysis was performed on bacteria recovered from protozoan food vacuoles at 2, 4 and 24 h post-feeding and compared with bacteria incubated under identical conditions in the absence of ciliates (control). Three biological replicates were analyzed for each condition at 2 h, and two biological replicates for each condition at 4 and 24 h. **(B)** Principal component analysis (PCA) showing that samples of intracellular bacteria at 2 h (blue) clustered distinctly from control samples at 2 h (grey) along principal component 1 (PC1), indicating significant differences in their expression profiles. Each point represents a biological replicate. **(C-D)** Venn diagrams showing the number of upregulated (UP, log2FC ≥ 1, FDR-adjusted p-value < 0.05) and downregulated genes (DOWN, log2FC ≥ -1, FDR-adjusted p-value < 0.05) that are unique or shared across the three time points (2, 4 and 24 h). **(E-G)** Functional changes in *B. cenocepacia* K56-2 transcriptome based on COG classification. The blue and red bars indicate the percentage of genes of each functional category upregulated or downregulated, respectively, in *B. cenocepacia* encased within *T. elliotti* food vacuoles versus free bacteria incubated in Tris-HCl buffer for 2 h (E), 4 h (F) and 24 h (G). The category “function unknown” (COG category S) includes genes of unknown function in addition to those without an associated COG category. The *P* value of each category is shown (*** *P* < 0.001, ** *P* < 0.01, * *P* < 0.05). **(H-M)** GO biological process enrichment analysis of DEG. Enriched GO biological processes among upregulated (H-J) and downregulated (K-N) genes at 2, 4 and 24 h. Bars represent the number of DEG sequences associated with each GO term, and bar colors indicate the corresponding *P value* according to the scale shown in each panel.

The classification of DEG in functional categories according to the Cluster of Orthologous Groups (COG) database revealed that upregulated genes across all time points were significantly enriched in functions related to energy production, amino acid metabolism, coenzyme transport/metabolism, translation/ribosome biogenesis, cell envelope biogenesis, and post-translational modification/protein turnover/chaperones (Fig. 1 E-G). In contrast, downregulated genes were predominantly assigned to cell motility, DNA replication/recombination/repair and signal transduction mechanisms. This suggests that bacteria remodeled their metabolism and cell envelope during transit through protozoan phagosomes. Consistently, functional enrichment analysis using the Gene Ontology (GO) database showed that upregulated genes at 2 h were mainly involved in translation, energy generation, amino acid metabolism, oxidative stress response, protein folding and cell envelope organization (Fig. 1 H). Downregulated genes at 2 h were mainly involved in transcription regulation, signal transduction, chemotaxis and flagellar apparatus and iron uptake (Fig. 1 K). Translation, protein folding, oxidative stress response, and cell envelope reorganization remained among the most significantly enriched GO terms at 4 and 24 h, suggesting an adaptive response during intracellular transit (Fig. 1 I-J).

### Stress responses activated in *B. cenocepacia* during passage through *T. elliotti*

Several stress response genes were significantly overexpressed in *B. cenocepacia* during passage through *T. elliotti,* reflecting the harmful conditions encountered within protozoan phagosomes (Fig. 2 and Table S6A-D). Oxidative stress response genes included those encoding a variety of ROS- and RNS-scavenging enzymes (catalases and peroxidases, superoxide dismutases and nitric oxide digoxygenase), redoxins and their reductases, glutathione metabolism enzymes, systems for repairing oxidized proteins and Fe-S clusters (methionine sulfoxide reductases and the iron-<u>s</u>ulfur <u>c</u>luster system isc), and limiting membrane lipid peroxidation (Fig. 2 A and Table S6A). This suggests a multifaceted response to counteract oxidative radicals generated in the phagosome. Genes encoding the transcriptional regulators OxyR, SoxR and OhrR were downregulated, whereas those encoding the tRNA methyltransferases TrmB and TrmJ, which enhance translation efficiency of *katB* mRNAs in bacteria [34,35], were overexpressed. In agreement with our previous study [7], genes of the SOS response and DNA repair pathways (Fig. 2 B and Table S6B) were upregulated, suggesting exposure to DNA damaging agents, most likely resulting from the oxidative burst in the phagosome. Because ROS primarily damages guanine to 8-oxoguanine (8-oxoG) [36], overexpression of genes involved in 8-oxoG detoxification and DNA repair (*mutM*, *mutY*, *mutT*, and 8-oxoG deaminase) suggests active repair of oxidative DNA/RNA lesions during passage through protozoan phagosomes. Genes encoding enzymes for the synthesis of compatible solutes such as the glycine-betaine (*betIBA* operon and additional betaine-aldehyde dehydrogenases) and trehalose (*otsA*, *otsB*), suggesting adaptation to phagosomal osmotic stress. In contrast, genes involved in glycine-betaine catabolism (*gbcAB*, *dgcAB, soxBDAG*) were repressed, suggesting that *B. cenocepacia* accumulated glycine-betaine during transit through protozoan phagosomes (Fig. 2 C and Table S6C).

**Fig. 2.**
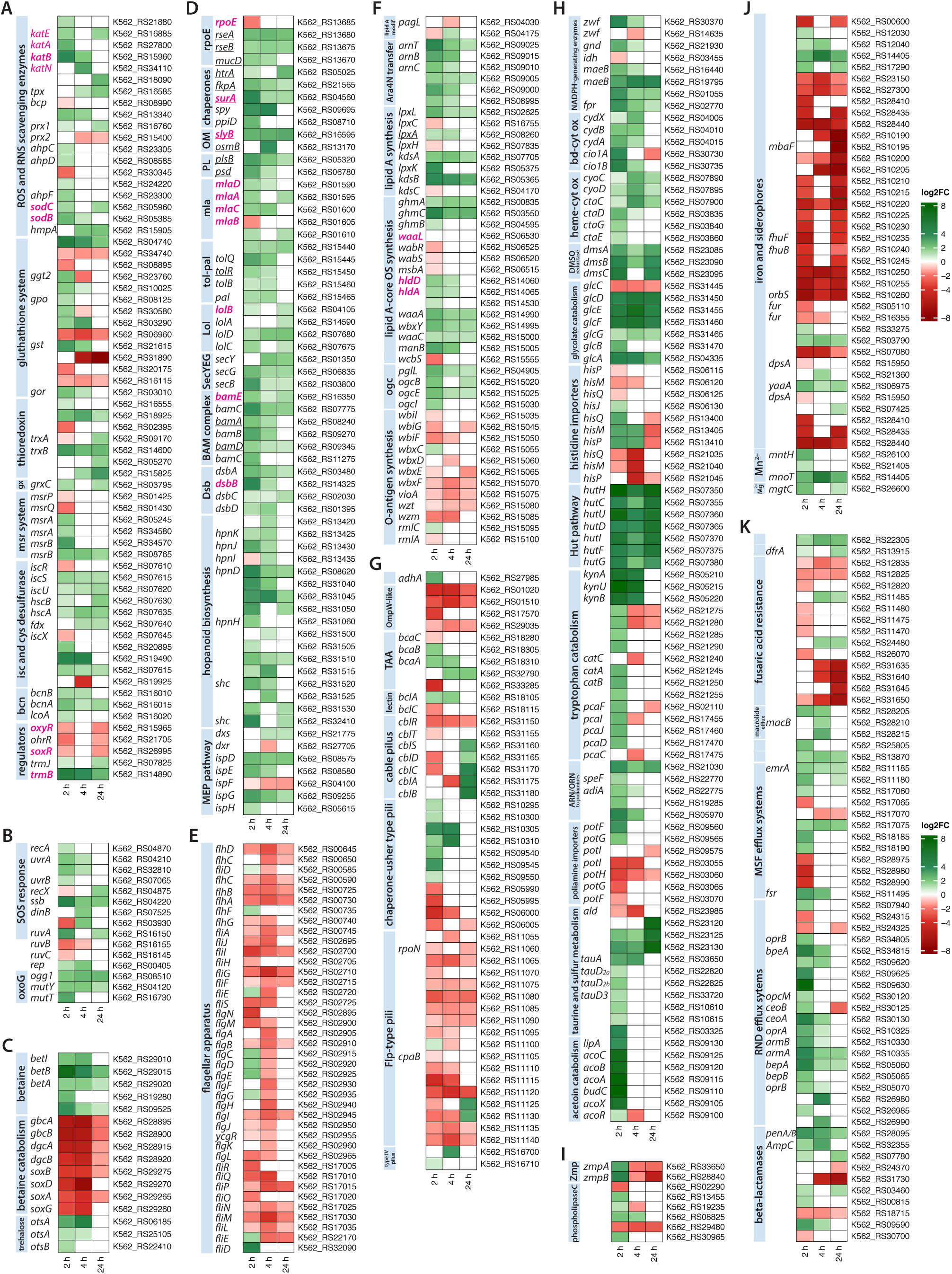
Expression kinetics of selected DEG during transit through protozoan food vacuoles. Heatmaps show the log2-fold change (log2FC) of genes associated with oxidative stress response (A), SOS response and DNA repair (B), osmotic stress (C), envelope stress response and outer membrane integrity and biogenesis (D), flagellar apparatus (E), LPS biosynthesis (F), adhesion (G), metabolic activities (H), proteases and phospholipases (I), iron and siderophore production (J) and antibiotic resistance (K). Each square represents the log2FC at 2, 4 and 24 h as indicated by the color scale. Assigned gene names are shown on the left, with corresponding locus tags on the right. Pink labels denote genes for which mutant strains were generated. Underlined names denote genes that have been reported to be part of the RpoE regulon in Gram-negative bacteria. Significance is based on FDR-padj < 0.05. ARN: arginine; BAM: beta-barrel assembly machinery; bcn: bacterial lipocalins; bd-cyt ox: cytochrome bd-type oxidase; Dsb: disulfide bond formation system; gx: glutaredoxin; heme-cyt ox: cytochrome heme-copper oxidase; Hut: histidine utilization pathway; isc: iron sulfur cluster; lol: localization of lipoproteins pathway; mla: mainteinance of lipid asymmetry pathway; MEP: non-mevalonate pathway; modif: modification; MSF: major facilitator superfamily; ogc: o-glycosylation gene cluster; OM: outer membrane: ORN: ornithine; OS: oligosaccharide; PL: phospholipid biosynthesis; RND: resistance-nodulation-division; Zmp: Zinc metaloprotease.

Upregulated genes associated with cell envelope integrity and biogenesis included the *rpoE* operon, which encodes the alternative sigma factor RpoE, a master regulator of the <u>e</u>nvelope <u>s</u>tress response (ESR) [37]; the *mlaFEDACB* cluster that <u>m</u>aintains outer membrane lipid <u>a</u>symmetry (Mla) [38]; lipoproteins that stabilize the OM during stress conditions (SlyB, Pal, BamD and OsmB) [39–42]; and periplasmic chaperones and proteases (HtrA, SurA, Spy) (Fig. 2 D and Table S6D). Furthermore, the protein glycosylation system [43] and major cell envelope assembly machineries, namely Lol (localization <u>o</u>f lipoproteins), SecYEG (general secretory pathway) and the BAM (β-<u>b</u>arrel <u>a</u>ssembly <u>m</u>achinery) complex were significantly induced, indicating lipoprotein trafficking, protein translocation across the OM and OMP assembly, respectively. Activation of the RpoE-mediated ESR and enhanced production of periplasmic chaperones and proteases points to OM damage and accumulation of unfolded proteins in the periplasm. Consistently, genes encoding the periplasmic <u>d</u>isulfide <u>b</u>ond formation <u>s</u>ystem (*dsbA, dsbB, dsbC, dsbD*), which aids in folding and stability of cell envelope proteins [44], were upregulated in protozoan phagosomes as well (Fig. 2 D and Table S6D). Since the Mla system helps to maintain OM barrier [38], its upregulation suggests a bacterial strategy to make the OM less permeable to harmful substances produced in the protozoan phagosome. In line with this, bacteria overexpressed genes involved in the biosynthesis of hopanoids (Fig. 2 D and Table S6D), which have been shown to contribute to membrane stability under low pH and membrane-damaging antibiotics in *B. cenocepacia* [45]. Consistently, bacteria also overexpressed genes encoding enzymes of the methylerythritol phosphate (MEP) pathway (Fig. 2 D and Table S6D) that in *Burkholderia* supplies isoprenoid precursors for hopanoid biosynthesis [46]. Additional changes in the bacterial OM included differential expression of genes associated with LPS synthesis (Fig. 2 F and Table S6F). While most genes involved in glycosylation of lipid A phosphate groups with L-Ara4N and lipid A-core oligosaccharide synthesis were upregulated during passage through protozoan phagosomes (at 2 and 4 h), O-antigen synthesis genes were downregulated, pointing to the production of rough LPS (lipid A + core) instead of smooth LPS.

### *B. cenocepacia* rewires bacterial metabolism to survive within protozoan phagosomes

Beyond upregulating stress response genes, our RNA-seq data suggest that *B. cenocepacia* underwent a dramatic metabolic reprogramming to exploit available resources and to cope with stress faced inside protozoan phagosomes. Genes encoding enzymes involved in NADHP production (Fig. 2 H and Table S6H) were upregulated likely to sustain the elevated NADPH demand imposed by antioxidant defenses. At 2 and 4 h, the two bd-type cytochrome oxidases (*cydABX* and *cio1AB*) exhibited higher expression than heme-type cytochrome oxidases (*ctaCDGE*, *cyoABCD*) suggesting limited oxygen availability within protozoan food vacuoles. In line with this, upregulation of terminal reductases for dimethylsulfoxide (*dmsABC*) points to the use of alternative electron acceptors under the low-oxygen tension in the phagosome. Alongside central carbohydrate metabolism pathways, bacteria upregulated genes associated with assimilation of diverse carbon sources such as glycolate, taurine, acetoin and different amino acids. The coordinated induction of glycolate permease *glcA* and the *glcDEFGB* operon (which converts glycolate into malate for entry into the TCA cycle) points to glycolate availability within protozoan food vacuoles, consistent with its role as inducer of *glcDEFGB* transcription [47]. Upregulation of amino acid uptake and metabolism suggests a strategy to exploit host amino acids as sources of carbon and nitrogen. Among these, the histidine utilization (Hut) pathway, which catabolizes histidine to glutamic acid [48], stood out, with the *hutHCUDIFG* (K562_RS07350-RS07380) cluster ranking among the top 20 upregulated genes in the dataset (Fig. 2 H and Tables S5 and S6H). Likewise, genes of the kynureine pathway (the *kynBUA* operon and cluster K562_RS21275-RS21290) were strongly induced. This pathway converts tryptophan to anthranilate and subsequently to catechol, which enters the beta-ketoadipate pathway to replenish intermediates in the TCA cycle [49]. Genes involved in arginine/ornithine conversion to polyamines and their importers were overexpressed as well (Fig. 2 H and Table S6H), indicating a potential strategy to scavenge host-derived polyamines. Upregulation of genes associated with taurine uptake and degradation to acetyl-CoA and sulfite may reflect that K56-2 exploits protozoan-derived taurine as a source of both carbon and sulfur (Fig. 2 H and Table S6H). Transport systems for sulfate and thiosulfate were also strongly upregulated, together with enzymes channeling sulfur into L-cysteine biosynthesis (Fig. 2 H and Table S6H), indicating a broader enhancement of sulfur acquisition and assimilation.

However, genes involved in siderophore-mediated iron acquisition were significantly downregulated during transit through protozoan phagosomes, suggesting that *B. cenocepacia* was not deprived from iron (Fe^2+^) within *Tetrahymena* food vacuoles (Fig. 2 J and Table S6J). This shift was accompanied by repression of the alternative sigma factor *orbS* and the ferric uptake regulator *fur*, consistent with their well-known repression under iron-replete conditions [50,51]. Downregulation of iron uptake genes occurred concurrently with upregulation of *yaaA* (K562_RS06795), which lowers intracellular Fe^2+^ levels to limit peroxide buildup under H_2_O_2_ stress [52], pointing to a coordinated strategy to minimize ROS generation via Fenton reactions. In contrast, genes encoding manganese import systems (*mntH*, K562_RS21405, K562_RS14405) were overexpressed at 2 h (Fig. 2 J and Table S6J). K562_RS14405 is homologous to MnoT (BTH_I1598), a TonB-dependent outer membrane receptor involved in Mn^2+^ scavenging in *B. pseudomallei* [53], suggesting an increased demand for Mn^2+^ in phagosome-trapped bacteria, consistent with its role as a cofactor in antioxidant enzymes [54]. Likewise, *mgtC*, which encodes a protein required for growth under magnesium-limiting conditions and intramacrophage survival in *B. cenocepacia* [55] and other bacteria [56] was overexpressed in protozoan phagosomes as well, pointing to adaptation to Mg^2+^ limitation within this intracellular niche.

### Passage through protozoan phagosomes enhances bacterial adhesion to human epithelial cells and resistance towards membrane-targeting antimicrobials

Several genes associated with host-cell attachment were upregulated during transit through protozoan phagosomes (2-4 h), such as the cable pilus associated adhesin *adhA*, the trimeric autotransporter adhesins (TAA) *bcaA*, *bcaB* and K562_RS32790; lectin *bclA*; and two chaperone-usher-type fimbria clusters (K562_RS10295-RS10310, K562_RS09540-RS09550) (Fig. 2 G and Table S6G). Moreover, bacteria packaged within protozoan EFVs (24 h) overexpressed the cable pilus operon *cblBACDS*, a specific virulence factor of the ET12 lineage [57]. Furthermore, genes encoding zinc metalloproteases ZmpA and ZmpB [58] and several phospholipases C (K562_RS13455, K562_RS19235, K562_RS08825, K562_RS30965) were overexpressed at 2 h (Fig. 2 I and Table S6I). Collectively, these results suggest that bacteria packaged in EFVs may have increased their capacity to adhere and invade host cells.

In contrast, genes associated with chemotaxis signaling and the flagellar apparatus were significantly downregulated in protozoan phagosomes (Fig. 2 E and Table S6E), potentially reflecting an energy-saving strategy to support stress responses and survival, or an adaptive mechanism to reduce host recognition. Consistently, the master regulator *flhDC* and the flagellar alternative sigma factor *fliA* [59] were downregulated as well.

Genes encoding components of the different secretion systems (TSS) T2SS, T3SS, T4SS, T6SS in *B. cenocepacia* were either downregulated or no differentially expressed during passage through *T. elliotti* (Table S6L), suggesting that they were not essential for survival in *T. elliotti* phagosomes. In contrast, transit through *T. elliotti* phagosomes dramatically upregulated genes linked to antibiotic resistance, primarily several RND-type efflux pumps, MSF efflux pumps, and the beta-lactamases PenB (the major beta-lactamase in *B. cenocepacia*) and AmpC (Fig. 2 K and Table S6K), which might enhance resistance towards antibiotics and other antimicrobials of EFV-packaged bacteria. Because the transcriptional profile of expelled bacteria suggested reduced OM permeability and enhanced adhesion potential, we next assessed their susceptibility to membrane-targeting antimicrobials as well as their adhesion capacity to human lung epithelial cells (A549 cell line) and collagen I, a component of epithelial extracellular matrix. Bacteria released from EFVs exhibited increased resistance to 10-30 μg/ml chlorhexidine compared to bacteria incubated in the absence of ciliates (*P* < 0.0001) (Fig. 3 A), consistent with OM impermeabilization or upregulation of efflux pumps during transit through protozoan phagosomes. Furthermore, in line with the overexpression of adhesin *bcaA,* which has been shown to bind to both collagen and A549 cells [60], bacteria expelled by ciliates exhibited higher adhesion to collagen-I-coated plates (*P* = 0.0345) and a modest, albeit non-significant (*P* = 0.12), increase in adhesion to A549 epithelial cells (Fig. 3 B-C).

**Fig. 3.**
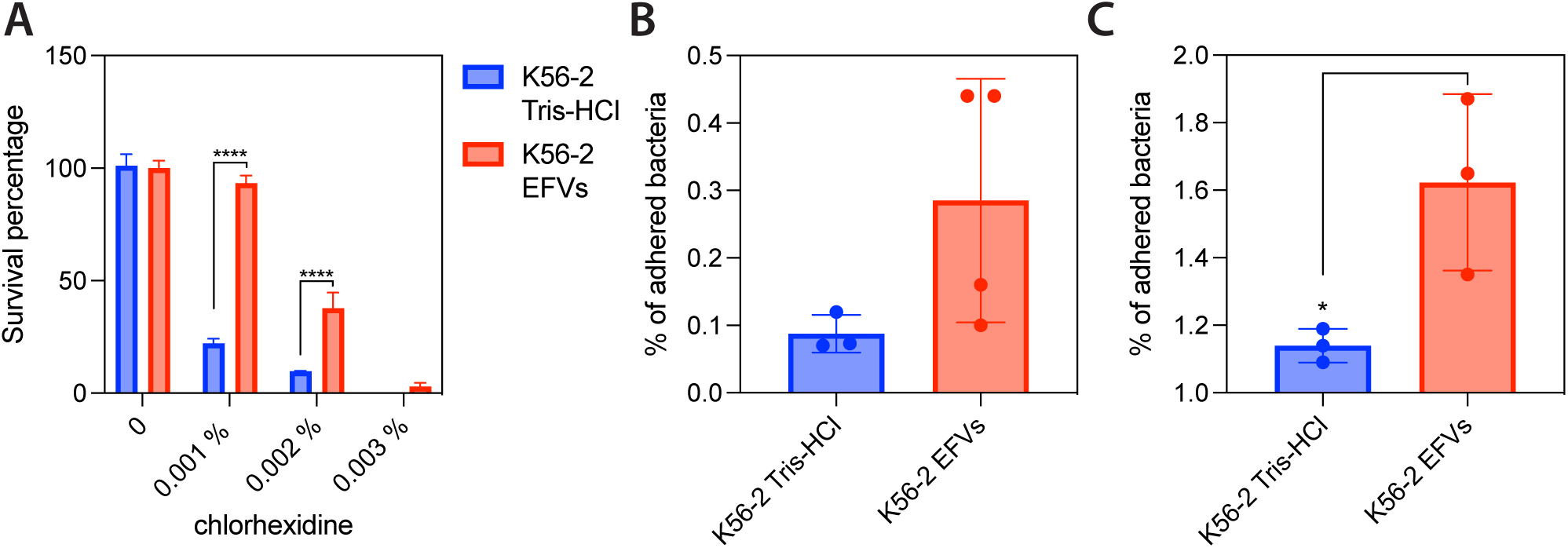
Passage through protozoa enhances bacterial adhesion capacity and resistance towards membrane-targeting antimicrobials. **(A)** Survival rate of K56-2 bacteria released from EFVs (red bars) and control bacteria (blue bars) exposed to chlorhexidine for 24 h in sterile saline solution. Survival is expressed as the percentage of viable bacteria relative to the initial bacterial count. Data represent the mean ± standard deviation of three independent biological replicates. Significant differences between EFV-released and control bacteria were determined using two-way ANOVA with Sidak’s multiple comparisons test (**** *P* < 0.0001). **(B-C)** Adhesion of K56-2 bacteria released from EFVs (red bar) and bacteria incubated in the absence of ciliates (control, blue bar) to A549 cells (A) or collagen-I-coated plates (B). Adhesion is expressed as the percentage of bacteria adhering relatively to the initial bacterial load added to the A549 monolayer or collagen-coated plates. Data represent the mean ± standard deviation of three independent biological replicates. Significant differences (* *P* < 0.05) between EFV-released and control bacteria were determined using unpaired Student’s t-test.

### Disruption of oxidative stress defenses and OM integrity dramatically impair *B. cenocepacia* intracellular survival in protozoa and macrophages

To assess whether *B. cenocepacia* employs similar strategies to survive phagocytosis by protists and macrophages, we generated mutant strains lacking components of *B. cenocepacia* stress responses and metabolic pathways upregulated during transit through *T. elliotti* phagosomes, and tested them for intracellular survival and growth within ciliates and THP-1 macrophages. Except the Δ*lolB* strain, all the tested mutants exhibited a growth rate in LB medium comparable to that of the isogenic parental strain K56-2 (Fig. S2). Regarding the oxidative stress response, deletion of *katB* severely impaired *B. cenocepacia* intracellular survival in *T. elliotti* (*P* = 0.0008) and THP-1 macrophages (*P* = 0.0031) (Fig. 4 A and C), whereas single deletion of each of the other catalases (*katA*, *katN*, K562_RS18090, K562_RS21880) had no effect (data not shown), underscoring the non-redundant role of KatB in protection against phagosomal oxidative radicals. Consistently, in parallel experiments only the Δ*katB* mutant displayed increased susceptibility (*P* < 0.01) to 4.26 mM H_2_O_2_ in LB compared to the isogenic wild type strain K56-2 (Fig. S3). Likewise, loss of *trmB*, which is expected to decrease the abundance of catalase KatB and other Phe- and Asp-enriched proteins [34], impaired intracellular survival in both phagocytes (Fig. 4 A and C), although the defect was statistically significant (*P* = 0.0007) only in *T. elliotti*. Deletion of both *sodB* and *sodC* superoxide dismutases significantly reduced bacterial survival in macrophages (*P* = 0.0083) but not in *T. elliotti* (Fig. 4 A and C), suggesting greater superoxide production in macrophage phagosomes compared to protozoan phagosomes. Deletion of *oxyR* reduced survival only in macrophages (*P* = 0.0021), while *soxR* was dispensable in both phagocytes (Fig. 4 A and C), consistent with the repression of *oxyR* and *soxR* during transit through *T. elliotti* phagosomes observed in our transcriptome data (Fig. 2, Table S6A) as well as in our previous study [7]. In line with this, deletion of *oxyR* or *soxR* did not reduce *B. cenocepacia* survival when exposed to 4.26 mM H_2_O_2_ in LB (Fig. S3). Besides, in agreement with this, RT-qPCR experiments revealed that the Δ*oxyR* strain retained full *katB* expression and, indeed, exhibited higher basal *katB* mRNA levels than the parental strain K56-2 (Fig. S4).

**Fig. 4.**
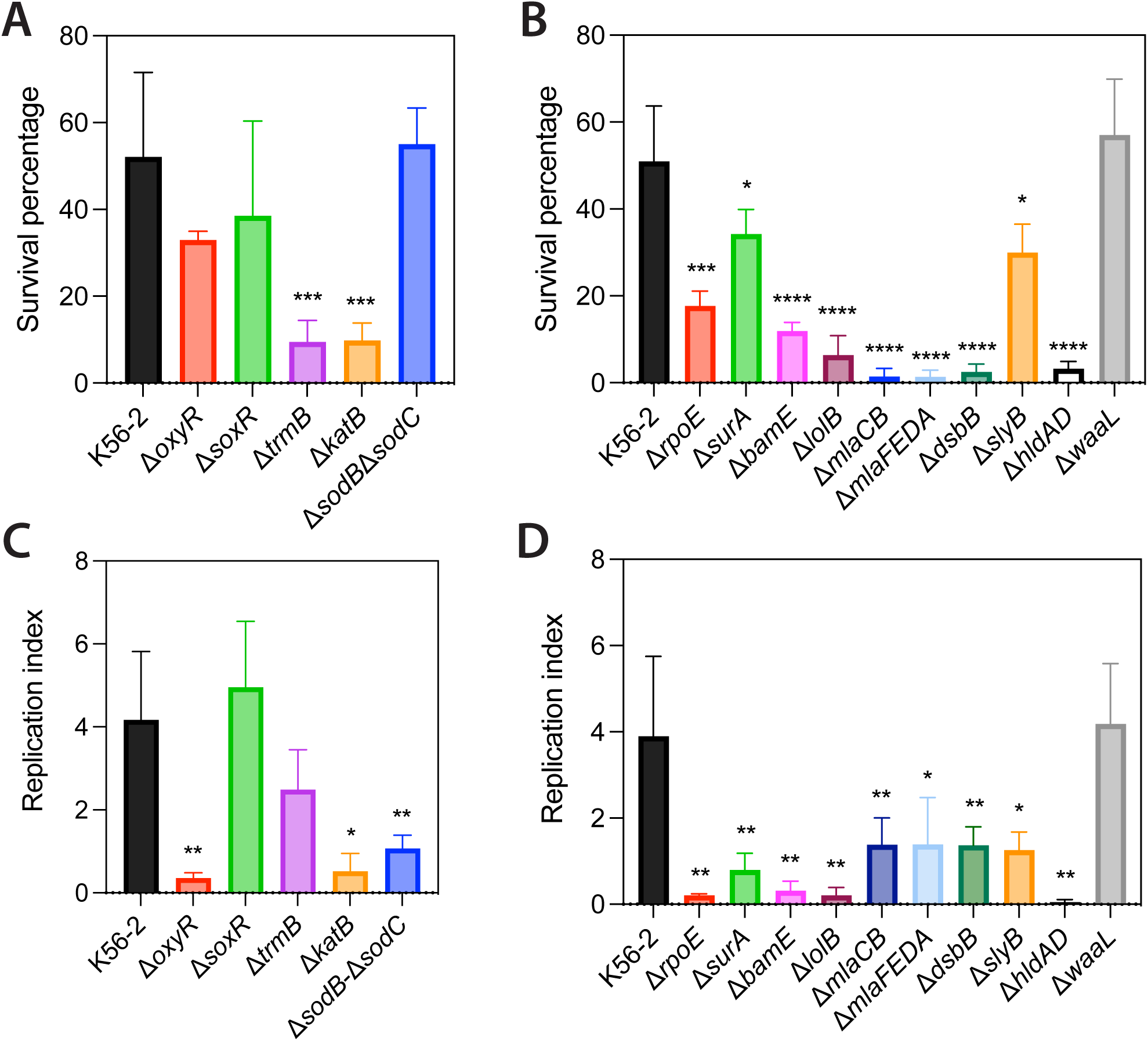
Intracellular survival of *B. cenocepacia* K56-2 and mutant strains in *T. elliotti* and THP-1 macrophages. **(A-B)** Intracellular survival in *T. elliotti* of K56-2 wild type (black bars) and mutants (colored bars) carrying deletions in genes involved in the oxidative stress response (A) and envelope stress or outer membrane integrity (B). Ciliates were incubated with bacteria (ratio 1:100 protist:bacteria) for 24 h int 0.1 M Tris-HCl buffer pH 7.5, lysed with 1% Triton X-100 and intracellular bacteria were enumerated by CFU counting. Survival was calculated as the percentage of recovered intracellular bacteria at 24 h relative to the initial bacterial inoculum. **(C-D)** Intracellular survival in THP-1 macrophages of K56-2 wild type (black bars) and mutants (colored bars) carrying deletions in genes involved in the oxidative stress response (C) and envelope stress or outer membrane integrity (D). Macrophages were infected for 2 h (MOI=2), treated with ceftazidime and amikacin for 2 h to kill extracellular bacteria (4 hpi), and further incubated for 20 h in fresh RPMI-1640 medium supplemented with gentamicin. Infected cells were lysed with 0.5% Triton X-100 at 4 or 20 hpi, and intracellular bacteria were quantified by CFU counting. Intracellular replication was expressed as a replication index, calculated as the ratio of bacteria recovered at 20 hpi relative to those recovered at 4 hpi. Data represent the average of at least three independent biological replicates. Statistical significances (**** *P* < 0.0001, *** *P* < 0.001, ** *P* < 0.01, * *P* < 0.05) between K56-2 and each mutant strain were determined with the unpaired t Student’s test.

Disruption of histidine utilization (Δ*hutHCUDIFG*) and kynurenine (Δ*kynBUA*) catabolic pathways did not impair intracellular survival within protozoan and macrophage phagosomes (data not shown), indicating that these metabolic pathways are dispensable under the conditions tested. In contrast, loss of *rpoE* and genes involved in OM integrity (*mlaCB, mlaFEDA, dsbB, lolB, slyB, bamE, surA*) caused significant (*P* < 0.05) intracellular survival defects in both phagocyte models (Fig. 4 B and D). Using the NPN uptake assay we found that all these mutants exhibited increased OM permeability compared to the parental K56-2 strain (Fig. 5). These results support a role for OM integrity in *B. cenocepacia* survival within phagocytes. Deletion of *hldAD*, which truncates the LPS core, significantly compromised bacterial survival in both phagocyte models, whereas loss of O-antigen ligase *waaL* and thus disruption of O-antigen synthesis, had no effect (Fig. 4 B and D), suggesting that an intact LPS core, but not the O-antigen, is required for *B. cenocepacia* intraphagosomal survival. Consistent with this result, Δ*hldAD* had increased OM permeability (*P* = 0.0027) whereas Δ*waaL* displayed OM permeability comparable to that of the parental strain K56-2 in the NPN assay (Fig. 5). Taken together, these results identify the ESR and maintenance of OM barrier function as major determinants of *B. cenocepacia* survival within phagocytes.

**Fig. 5.**
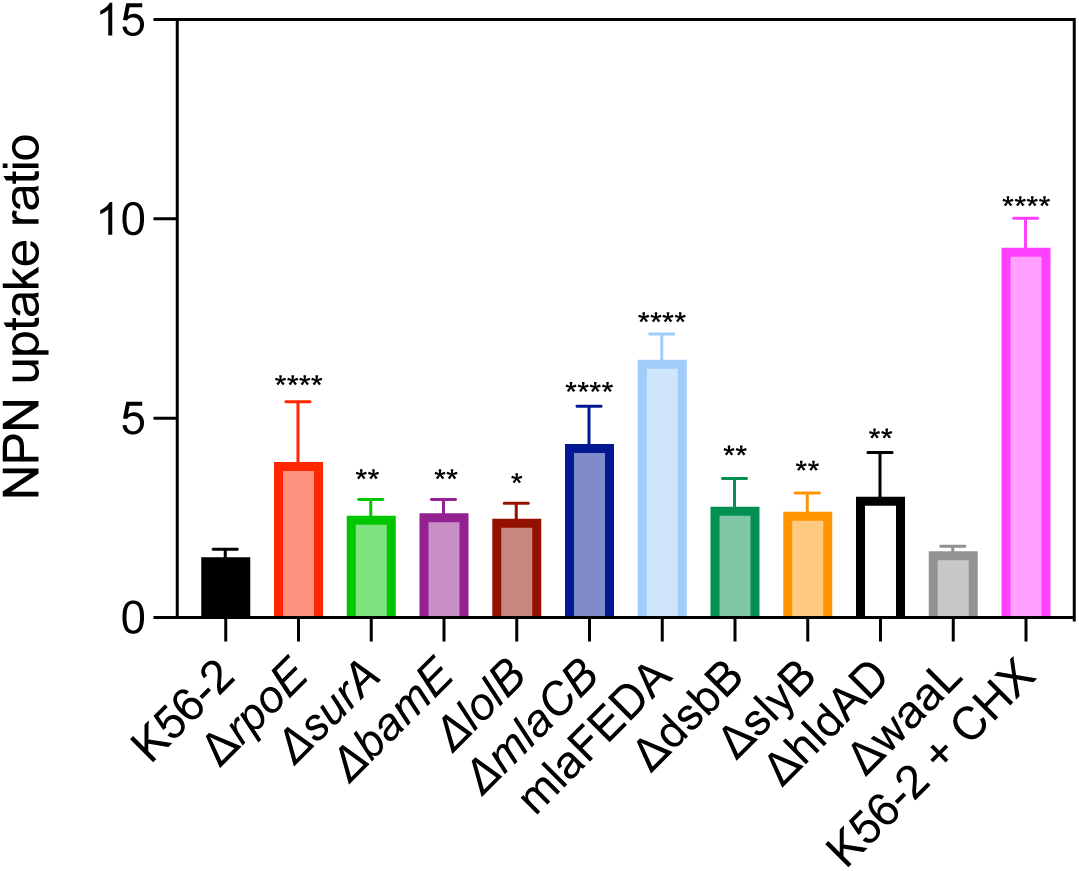
Outer membrane permeability of K56-2 wild type and mutant strains as measured by the NPN uptake assay. Synchronized (OD_600_ = 1) bacterial cultures were washed and incubated with 80 μM NPN in 5 mM HEPES buffer and 420-440 nm fluorescence was measured. For each strain the NPN uptake ratio was calculated as the net bacterial fluorescence (corrected for cell autofluorescence) divided by net buffer fluorescence (corrected for buffer background). Higher NPN uptake ratio indicates reduced OM integrity. K56-2 treated with 30 μg/ml chlorhexidine served as positive control for membrane permeabilization. Data show means of at least three independent experiments. Significant differences relative to wild-type K56-2 was determined using a one-way ANOVA with Dunnett’s multiple comparison test (**** *P* < 0.0001, *** *P* < 0.001, ** *P* < 0.01, * *P* < 0.05).

## Discussion

This study shows that, in order to survive phagocytosis by ciliates, *B. cenocepacia* activates a coordinated network of metabolic pathways and stress responses that allow the bacterium to exploit host-derived nutrients, ameliorate stress, repair damage and maintain redox balance. Importantly, stress responses triggered during transit through protozoan phagosomes not only facilitated intracellular survival but also simultaneously primed *B. cenocepacia* for enhanced resistance toward harsh conditions and antimicrobials, in agreement with our previous study [7]. Furthermore, passage through ciliates upregulated genes associated with host-cell adhesion, and expelled bacteria exhibited increased adherence to collagen and human lung epithelial cells indeed, underscoring the role of EFVs as vehicles for pathogen dissemination. Alongside energy-saving measures such as downregulation of the flagellar apparatus, bacteria redirected metabolic flux toward alternative carbon sources, while upregulating glutathione metabolism, cysteine biosynthesis and NADPH-generating pathways likely to sustain some of the antioxidant defenses. Non-flagellated *P. putida* cells display an increased NADPH/NADP^+^ ratio and higher oxidative stress tolerance than the wild type [61]. Thus, flagellar repression during transit through protozoan phagosomes might be a strategy to conserve reducing power for oxidative stress defenses. Furthermore, bacteria repressed genes linked to iron uptake, likely to limit additional ROS production via Fenton reactions, and upregulated genes involved in repair of oxidized proteins, DNA/RNA and lipids. Disruption of OM barrier function and oxidative defenses dramatically compromised intracellular survival, indicating that persistence within protozoan phagosomes requires the coordinated activation of antioxidant systems and mechanisms that preserve envelope integrity. Among the ROS-scavenging enzymes tested, catalase KatB was uniquely required for survival within ciliates and macrophages, consistent with its role as the major catalase of *B. cenocepacia* [62]. The survival defect of the Δ*trmB* mutant further highlights the importance of maintaining high levels of KatB for protection against phagosomal ROS, as TrmB positively modulates *katB* mRNA translation in bacteria [34]. The requirement for *oxyR*, *sodB* and *sodC* for survival in THP-1 but not in *T. elliotti,* suggests that macrophage phagosomes impose a stronger oxidative stress than Tetrahymena food vacuoles. The lack of phenotype for the Δ*soxR* mutant in both phagocytes suggests that, unlike enteric bacteria [63], *B. cenocepacia* does not rely on SoxR for protection against superoxide stress. This agrees with studies in *B. multivorans* and *P. aeruginosa* showing that SoxR primarily responds to redox-active metabolites such as phenazines rather than controlling canonical ROS detoxification pathways [64].

The Gram-negative OM acts as a highly effective permeability barrier against a variety of antimicrobials, including ROS. OM permeability barrier requires lipid asymmetry, which is maintained by the Mla system [38]. Our findings of impaired intracellular survival in evolutionarily distant phagocytes (protists and macrophages) for mutants in the Mla pathway confirm that this system not only preservers OM asymmetry but also underpins multiple fitness traits required for survival under diverse environmental and host-imposed stresses, consistent with recent studies linking it to bacterial resistance to antibiotics and human serum complement [65,66]. Therefore, the pleiotropic effects of inhibiting the Mla system highlight this pathway as a potential drug target to treat *Bcc* infections. In addition to the asymmetric nature of the OM, the LPS of *Burkholderia* species contains unique structural modifications that further reduce OM permeability such as D-*glycero*-α-D-*talo*-octulosonic acid (Ko) residues instead of 3-deoxy-D-*manno*-octulosonic acid (Kdo) in the LPS core oligosaccharide [67], and constitutive functionalization of lipid A and core oligosaccharide with 4-amino-4-deoxy-L-arabinose (Ara4N) residues [68,69]. Indeed, compared to *E. coli*, the OM of *Burkholderia* species is approximately 89% less permeable [70]. Although lipid A glycosylation with Ara4N is an essential process in *Burkholderia* species [68], its upregulation during transit through protozoan phagosomes might indicate OM reinforcement to resist phagosome-imposed stress. The LPS core oligosaccharide contributes to the OM barrier function by restricting the passage of hydrophobic molecules, while the acyl chains within the bilayer also help limit the entry of many hydrophilic compounds [71]. Our work shows that a complete LPS core is required for intracellular survival in both protozoan and macrophage phagosomes, while the O-antigen seems to be dispensable, as disruption of O-antigen synthesis did not impair bacterial intraphagosomal survival, and O-antigen synthesis genes were indeed repressed during transit through protozoan phagosomes. In line with our results, previous studies have shown that a complete LPS core rather than the O-antigen is required for OM integrity in *B. cenocepacia* [66], and that loss of polymeric O-antigen does not increase sensitivity to antimicrobial peptides [72,73], which are expected to be produced in *Tetrahymena* phagosomes as well [74]. Importantly, it has been recently proposed that production of rough LPS helps enterobacteria to enhance OM integrity [75]. Therefore, our results suggest that repression of O-antigen synthesis may represent a survival strategy by strengthening OM permeability barrier and enhancing resistance to the harsh conditions encountered in the protozoan phagosome. In line with our hypothesis, adaptive mutations in *rfbP* were recently identified in *Salmonella* populations under predation by *Tetrahymena*, resulting in a rough LPS phenotype that enhanced grazing resistance [76]. However, while Huang and coworkers suggested that O-antigen deficiency enabled *Salmonella* to evade recognition by the *Tetrahymena* predator, such a mechanism contrasts with the established feeding behavior of *Tetrahymena* species, which primarily filter prey based on size rather than cell-cell contact recognition [77]. Consistent with earlier work [78], we have found no feeding preferences in *T. elliotti* based on the presence or absence of O-antigen in *B. cenocepacia*, even though these variations can be readily distinguished by amoebae [78]. O-antigen loss did not prevent bacteria ingestion by *T. elliotti*. Therefore, we propose that the rough LPS phenotype does not function as an evasion strategy, but instead may contribute to *B. cenocepacia* survival by strengthening OM integrity.

The integrity of the bacterial cell envelope is monitored and maintained by the envelope stress response (ESR) [37]. *B. cenocepacia* genomes lack homologues to the Bae, Rcs and Cpx signaling pathways that regulate ESR in bacteria, thus only the RpoE-dependent ESR has been identified in this species [79]. Our results show that RpoE itself and putative RpoE-regulated genes such as *surA* and *bamE* are required for intracellular survival within protists and macrophages, in agreement with previous studies showing that RpoE is important for delaying phagolysosomal fusion in macrophages [79]. Both *surA* and *bamE* are part of the RpoE regulon in Gram-negatives [80–83]. Although their roles have not been investigated in *Burkholderia* species yet, it is well recognized that the BAM complex and associated shuttle proteins are critical for membrane integrity and stability [39,84]. SurA is the primary periplasmic chaperone in the OMP biogenesis network that delivers unfolded beta-barrel proteins to the BAM complex for folding and insertion into the OM [84]. Disruption of either protein alters OMP composition and impairs OM barrier function and therefore resistance to phagosomal antimicrobials [85,86]. SlyB has been associated with OM integrity in *B. multivorans* and other Gram-negative bacteria [42,87]. Consistent with our results, it has been reported that *slyB* transcription is dramatically induced by the stress response accompanying loss of OM asymmetry and low levels of Mg^2+^ via activation of the phoPQ two-component system in *E. coli* [42]. In *P. aeruginosa slyB* expression is regulated by RpoE [81]. Both conditions known to induce *slyB* expression (Mg^2+^ limitation and OM damage) are encountered within protozoan phagosomes according to our RNA-seq data.

Given that accumulation of unfolded OMPs and damaged LPS activates the rpoE-dependent pathway [88], it is tempting to speculate that RpoE activation under OM stress in *B. cenocepacia* might shift LPS biosynthesis toward the rough form by repressing O-antigen polymerization. This mode of LPS remodeling could be adaptive as it might help *B. cenocepacia* to strength OM permeability barrier during transit through the phagosome. Under RpoE-inducing conditions, *E. coli* alters LPS biosynthesis via RpoE-regulated sRNAs, particularly RybB, which represses *waaR* and leads to the truncation of the LPS outer core [89]. Although a RybB homolog has not been found yet in *B. cenocepacia*, RpoE activation might similarly drive post-transcriptional remodeling of LPS composition to survive intracellular digestion by protists. This hypothesis remains to be experimentally confirmed. The RpoE pathway is poorly characterized in *Burkholderia* species and research underway in our laboratory might help us to establish if RpoE drives LPS remodeling as part of *B. cenocepacia* strategy to withstand phagosomal conditions.

Overall, this study provides new insights into the molecular mechanisms that enable *B. cenocepacia* to resist phagosomal digestion in evolutionarily distant phagocytes (protozoa and macrophages), and identifies processes responsible for the enhanced stress tolerance and antibiotic resistance acquired following passage through protozoan phagosomes. Cell envelope remodeling emerged as a central adaptive strategy for intracellular survival but also provided bacteria with colonization advantages beyond the intracellular niche, including enhanced resistance to antimicrobials and increased adhesion to human airway epithelial cells. Besides, this study underscores the need to dissect the stress responses and regulatory networks that enable bacteria to survive phagocytosis by protists and to determine how these adaptive mechanisms contribute to bacterial persistence in both environmental reservoirs and humans, ultimately priming bacteria for infection.

Overall, our findings suggest that exposure to protozoan phagosomes induce adaptive responses that enhance bacterial fitness across phagocytic hosts, potentially linking environmental adaptation to survival in the human host.

## Acknowledgements

This study was supported by grants PID2020-113540GB-I00 and PDI2024-157739NB-I00 from the Spanish State Research Agency/Ministry of Science and Innovation awarded to F.A. A.M. was supported by research contract PEJ-2021-AI/SAL-21691 from the Madrid Government (NextGeneration EU funds). N. C. and I. B. were supported by research contracts funded through the Investigo Program (NextGeneration EU funds). We thank Dr. María Jesús Grilló (IDAB-CSIC, Spain) and Dr. Alicia Sánchez-Gorostiaga (iMiDRA, Spain) for kindly providing strains HB101/pRK2013 and BW2514, respectively. Plasmids pGPI-SceI and pDAI-SceI-SacB were a gift from Dr. Miguel Valvano (Addgene plasmids #32060 and #113635).

## Data availability statement

Any further information or additional data will be made available upon request.

