## Supplementary material for "Transcriptomics uncovers outer membrane remodeling as conserved strategy for *Burkholderia cenocepacia* survival in protozoan and macrophage phagosomes": Suplementary Tables S1-S4 and Figures S1-S4

Department of Genetics, Physiology and Microbiology. Faculty of Biological Sciences, Complutense University of Madrid. 28040 Madrid. Spain.

**Supplementary Tables**

**Table S1**. Bacterial strains used in this study.

| Strain | Description | Source |
|---|---|---|
| *E. coli* | | |
| HB101/pRK2013 | HB101 strain harbouring pRK2013 | M. J. Grillo (IDAB-CSIC) |
| BW25141 | F-, *Δ(araD-araB)567*, *ΔlacZ4787*(*::rrnB*-3), *Δ(phoB-phoR)580*, *$\lambda^{-}$*, *galU95*, *ΔuidA3::pir*$^{+}$, *recA1*, *endA9*(del-ins)*::FRT*, *rph-1*, *Δ(rhaD-rhaB)568*, *hsdR514* | A. Sánchez-Gorostiaga (iMiDRA, Spain) |
| *B. cenocepacia* | | |
| K56-2 | Clinical isolate from a CF patient, ET12 lineage | BCCM/LMG Bacterial Collection |
| Δ*oxyR* | Δ*oxyR* derivative of K56-2 | This study |
| Δsox*R* | Δ*soxR* derivative of K56-2 | This study |
| ΔtrmB | Δ*trmB* derivative of K56-2 | This study |
| Δ*katB* | Δ*katB* derivative of K56-2 | This study |
| Δ*sodB*Δ*sodC* | Δ*sodB*Δ*sodC* derivative of K56-2 | This study |
| Δ*rpoE* | Δ*rpoE* derivative of K56-2 | This study |
| Δ*surA* | Δ*surA* derivative of K56-2 | This study |
| Δ*bamE* | Δ*bamE* derivative of K56-2 | This study |
| Δ*lolB* | Δ*lolB* derivative of K56-2 | This study |
| Δ*mlaCB* | Δ*mlaCB* derivative of K56-2 | This study |
| Δ*mlaFEDA* | Δ*mlaFEDA* derivative of K56-2 | This study |
| Δ*dsbB* | Δ*dsbB* derivative of K56-2 | This study |
| Δ*slyB* | Δ*slyB* derivative of K56-2 | This study |
| Δ*hldAD* | Δ*hldAD* derivative of K56-2 | This study |
| Δ*waaL* | Δ*waaL* derivative of K56-2 | This study |
| Δ*katA* | Δ*katA* derivative of K56-2 | This study |
| Δ*katN* | Δ*katN* derivative of K56-2 | This study |
| ΔK562_RS10890 | ΔK562_RS10890 derivative of K56-2 | This study |
| Δ*hutHCUDIFG* | Δ*hutHCUDIFG* derivative of K56-2 | This study |
| Δ*kynBUA* | Δ*kynBUA* derivative of K56-2 | This study |

**Table S2**. Plasmids used in this study.

| Plasmid | Description | Reference |
|---|---|---|
| pRK2013 | ori$_{colE1}$, RK2 derivative, Kan$^r$ mob$^+$ tra$^+$ | (Figurski & Helinski, 1979) |
| pGPI-SceI | ori$_{R6K}$, mob$^+$,Tp$^r$, I-SceI-I restriction site | (Flannagan et al., 2008) |
| pDAI-SceI-SacB | ori$_{pBBR1}$, mob$^+$,Tet$^r$, expressing I-Sce-I and sacB | (Flannagan et al., 2008) |
| pIB022 | pGPI-SceI with regions flanking K56-2 *oxyR* | This study |
| pIB023 | pGPI-SceI with regions flanking K56-2 *soxR* | This study |
| pFA234 | pGPI-SceI with regions flanking K56-2 *trmB* | This study |
| pIB025 | pGPI-SceI with regions flanking K56-2 *katB* | This study |
| pEF001 | pGPI-SceI with regions flanking K56-2 *sodB* | This study |
| pEF002 | pGPI-SceI with regions flanking K56-2 *sodC* | This study |
| pAM027 | pGPI-SceI with regions flanking K56-2 *rpoE* | This study |
| pAM011 | pGPI-SceI with regions flanking K56-2 *surA* | This study |
| pAM018 | pGPI-SceI with regions flanking K56-2 *bamE* | This study |
| pAM020 | pGPI-SceI with regions flanking K56-2 *lolB* | This study |
| pFA283 | pGPI-SceI with regions flanking K56-2 *mlaCB* | This study |
| pFA284 | pGPI-SceI with regions flanking K56-2 *mlaFEDA* | This study |
| pFA235 | pGPI-SceI with regions flanking K56-2 *dsbB* | This study |
| pFA299 | pGPI-SceI with regions flanking K56-2 *slyB* | This study |
| pAM115 | pGPI-SceI with regions flanking K56-2 *hldAD* | This study |
| pFA315 | pGPI-SceI with regions flanking K56-2 *waaL* | This study |
| pIB024 | pGPI-SceI with regions flanking K56-2 *katA* | This study |
| pIB017 | pGPI-SceI with regions flanking K56-2 *katN* | This study |
| pIB028 | pGPI-SceI with regions flanking K56-2 K562_RS10890 | This study |
| pFA282 | pGPI-SceI with regions flanking K56-2 *hutHCUDIFG* | This study |
| pAT007 | pGPI-SceI with regions flanking K56-2 *kynBUA* | This study |

**Table S3**. Primers used in this study. Nucleotides introduced to create restriction enzyme sites are shown in lowercase letters.

| Primer name | Sequence (5' → 3') | Application |
|---|---|---|
| FA308oxyRUP1 | TTTtctagaTCGGTGTGCGTACTGCT | Generation of plasmids and mutant strains |
| FA309oxyRUP2 | TATatcgatGCTGGCTCACGAAGCAC | |
| FA310oxyRDS3 | TATatcgatGTATGGCGCAAGAGCTTC | |
| FA311oxyRDS4 | TTAgaattcCGTGTATCGGACATGCAAT | |
| FA314soxRUP1 | TTTtctagaGCGTTGTCGAGTTCTTCCT | |
| FA315soxRUP2 | TTTctcgagCCGGGAAAGCTTGACTTGA | |
| FA316soxRDS3 | TTTctcgagCCGACCGATTGCGAATGA | |
| FA317soxRDS4 | TTTgaattcCTCGACGCGAACCTGAAA | |
| FA603trmBUP1 | ATTtctagaCGAAGGACCAGTTCGTTCAA | |
| FA604trmBUP2 | ATTctcgagGTACAGAGAAAGGCTCGGTTC | |
| FA605trmBDS3 | ATTctcgagGGAACCGACGCTCGAAA | |
| FA606trmBDS4 | ATTgatatcCACGCTGAACGTCGTGAT | |
| FA326katBUP1 | TTTtctagaCTTGTCGCTGATACCGATGTT | |
| FA327katBUP2 | TTTctcgagGCCTGGAACAAGGTGATGAA | |
| FA328katBDS3 | TTTctcgagCGGCTTCTCCTGTCGTACT | |
| FA329katBDS4 | TTTgaattcCCGGCCACGATGAATTGAC | |
| FA658sodBUP1 | ATTtctagaCCGCACATTCCGGTGAT | |
| FA659sodBUP2 | ATTccatggTCCTTCGTTGAGTATGTAGGGA | |
| FA660sodBDS3 | ATTccatggACACGCGTACTACATCGACTA | |
| FA661sodBDS4 | ATTgaattcTGATGGCTTTGAGTCGTCTTG | |
| FA663-sodCUP1 | ATTtctagaATTCTCGACGCCGACATCTAC | |
| FA664-sodCUP2 | ATTccatggGGTAAAGGGACCCGGGAAA | |
| FA665-sodCDS3 | ATTccatggCGGCGTGATCCGTTGAC | |
| FA666-sodCDS4 | ATTgatatcCGGAAACTGGTGAGACGTTATG | |
| FA408rpoEUP1 | ATTtctagaAAGGTATTGGTCGAGGCTGGCG | |
| FA409rpoEUP2 | ATTcatatgGAAGGCAAGCGCTGGTAAGCG | |
| FA410rpoEDS3 | ATTcatatgTAACTAGGGATGCAACGCGGAG | |
| FA411rpoEDS4 | ATTgaattcTTGACGAGTCCATTCGATGCTCC | |
| FA295surAUP1 | ATTtctagaCTACTTCGCCGATCAGCGCGTCAC | |
| FA296surAUP2 | ATTccatggGCAGACATGCCGGCGTTACTCGT | |
| FA297surADS3 | ATTccatggGAACAGGCCTATGCGGACTGGCT | |
| FA298surADS4 | ATTgaattcTTCAGCGGCAAGTGGGTGGTCGCGA | |
| FA392bamEUP1 | ATTtctagaGATGATCTCGATGTCGTAGCCGT | |
| FA393bamEUP2 | ATTgtcgacCAAGGCTCGGACAACCTGCAG | |
| FA394bamEDS3 | ATTgtcgacGTACGACGAACAACCCGCCA | |
| FA395bamEDS4 | GAAgaattcCTCGACGCGGC | |
| FA396lolBUP1 | ATTtctagaAGTTGCCACAGGCGGTTCA | |
| FA397lolBUP2 | ATTccatggACGCGCGTCAAGCAGATCC | |
| FA398lolBDS3 | ATTccatggCATCAGAGGCCGTTGATCTGC | |
| FA399lolBDS4 | ATTgaattcATTGTCGACAAACAGGGCAAGACCGACGA | |
| FA711mlaCBUP1 | ATTtctagaCCGGACGATGTACAAGTTCAA | |
| FA712mlaCBUP2 | ATTccatggTCCGGCTGTAACGAATGTAAG | |
| FA713mlaCBDS3 | ATTccatggATTGATCCTGTCCGACCGA | |
| FA714mlaCBDS4 | ATTgaattcAACCGCAGATACAGTTGCAG | |
| FA707mlaFEDAUP1 | ATTtctagaCTCGCGGACGACTGGAT | |
| FA708mlaFEDA UP2 | ATTccatggGGCCTGCATTGGAATGATGT | |
| FA709mlaFEDA DS3 | ATTccatggCTGCACTGACGGACTTACAT | |
| FA710mlaFEDA DS4 | ATTgaattcGCGGACGAGTCGAACTG | |
| FA607dsbBUP1 | ATTtctagaCTCGTCGACCGGATGCT | |
| FA608dsbBUP2 | ATTccatggGTTCCTCGTCGGTCGGT | |
| FA609dsbBDS3 | ATTccatggGCGCGTAAACACCTACCC | |
| FA610dsbBDS4 | ATTgaattcCTTCGCGAGCGACATGAT | |

| | | |
|---|---|---|
| FA723slyBUP1 | ATTtctagaCTACGACTCCATCATGACCTT | Generation of plasmids and mutant strains |
| FA724slyBUP2 | ATTccatggTCATTTGAGACTGTAACCGAATG | |
| FA725slyBDS3 | ATTccatggTCGAACGCATGTGCGAA | |
| FA726slyBDS4 | ATTgaattcGAAGAAACCGGCTGCTAAGA | |
| FA635hldADUP1 | ATTtctagaCTTGCTGATCATCGACAGCGC | |
| FA636hldADUP2 | ATTcatatgTAAACCGGCGCGTTGCATC | |
| FA637hldADDS3 | ATTcatatgCGGAACGACTTCGCGAAGAG | |
| FA638hldADDS4 | ATTgatatcGAATCTCGCCGACAAGGTCG | |
| FA753waaLUP1 | ATTtctagaATCCAGGCGTTGGCTTT | |
| FA754waaLUP2 | ATTccatggTCATCGAACTTGCACCAGAA | |
| FA755waaLDS3 | ATTccatggTGAACGCCGTCGCGAGC | |
| FA756waaLDS4 | ATTgaattcCCGCAGTTCGTCCGCAA | |
| FA279katNUP1 | TTTtctagaCATCCGTACTCCTTCACGAAA | |
| FA280katNUP2 | TTActcgagACTGAAGCCGTGTGTTGT | |
| FA281katNDS3 | TTActcgagCGGTATTCCGGAAGGCAT | |
| FA282katNDS4 | TTAgaattcGCTGCACGTCGAACTTGA | |
| FA320katAUP1 | TTTtctagaACGCTGATGGTCAAGACAGAC | |
| FA321katAUP2 | TTTctcgagTATGAGGACGTGTGGGTGTA | |
| FA322katADS3 | TTTctcgagATTTCTTCCGCAACCTGCTC | |
| FA323katADS4 | TTTgaattcCGATTTCGATCGCGGTCATT | |
| FA366RS18090UP1 | TTAtctagaGCGCATCAGCTCGACATT | |
| FA367RS18090UP2 | TTTctcgagCGAATCGATTGAGGTACGAACG | |
| FA368RS18090DS3 | TTTctcgagCGAACGATCCGATCCTGAAC | |
| FA369RS18090DS4 | TTAgaattcCAGGTTGACGCCCTTGAA | |
| FA749BM0931UP1 | ATTtctagaGCCACGCTGCTGAAACA | |
| FA750BM0931UP2 | ATTccatggCGTCAGACGATTGGCTAATTGATA | |
| FA751BM0931DS3 | ATTccatggGCGATGCTGTCGTACTTCTAC | |
| FA752BM0931DS4 | ATTgaattcGCATTGTTCAGACGGTTGATTC | |
| FA453KynBUAUP1 | ATTtctagaTGTTCACCGATCGAAGGACG | |
| FA454KynBUAUP2 | ATTccatggAAGATGCTCGACGTCGTGCT | |
| FA455kynBUADS3 | ATTccatggAATGATAGGTGGACGCTCGCA | |
| FA456KynBUADS4 | ATTgaattcAAGGCGACCACGTGATCTTTG | |
| FA703BL2240-6UP1 | ATTtctagaCGTAGGCTTCCGACGATTG | |
| FA704BL2240-6UP2 | ATTccatggACTCGGGAGTTTCCCTGAT | |
| FA705BL2240-6DS3 | ATTccatggCCGATGCAACGCACAAA | |
| FA706BL2240-6DS4 | ATTgaattcAATAGCTGCCGGTCAACA | |
| gyrAFw | CGACGCAGATGAAGGAAGA | Quantitative RT-PCR |
| gyrARv | CATAGACCTTGACCCAGTACAC | |
| oxyRFw | TCGTCGCTCGAAACCATTC | |
| oxyRRv | GCACGTACGACAGCAGTT | |
| katBFw | AGTGGAGCAACGACTTCTTC | |
| katBRv | AATCACCTCGTCCGCATC | |

**Table S4.** Number of reads mapped to the *B. cenocepacia* K56-2 reference genome per sample.

| Condition | Sample ID | Number of quality control passed reads | Number of reads mapped to the K56-2 genome |
|---|---|---|---|
| Control 2 h | C4C2I | 40036456 | 39708246 |
| | C5C2G | 34371667 | 33936835 |
| | C6C2H | 41851816 | 41274477 |
| K56-2 + *T. elliotti* 2 h | T1T2A | 10027340 | 9330820 |
| | T2T2C | 10600637 | 9746843 |
| | T3T2F | 7514677 | 6601080 |
| Control 4 h | S5C410 | 20584891 | 20416260 |
| | S6C411 | 18170708 | 17979716 |
| K56-2 + *T. elliotti* 4h | S1T4E | 30247672 | 7375839 |
| | S2T4F | 5410433 | 901269 |
| Control 24 h | S7C241 | 15610286 | 15440695 |
| | S8C242 | 18513486 | 18094557 |
| K56-2 + *T. elliotti* 24 h | S3T24C | 10566461 | 6878556 |
| | S4T24D | 5461698 | 3533922 |

**Supplementary Tables S5 – S6** are provided as separate Excel files

**Table S5**. List of differentially expressed genes (DEG) in *B. cenocepacia* K56-2 during passage through *T. elliotti* at 2, 4 and 24 h.

**Table S6**. Selected DEG associated to stress responses and different pathways in *B. cenocepacia* K56-2 during passage through *T. elliotti* at 2, 4 and 24 h.

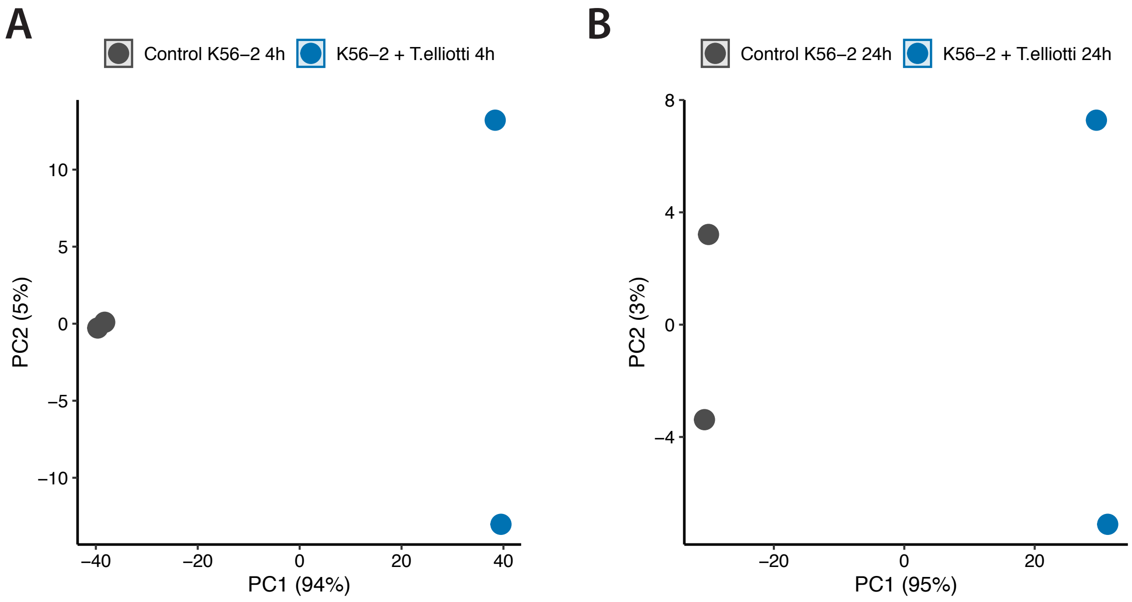

**Fig S1**. Principal component analysis (PCA) showing that samples of intracellular bacteria (blue) clustered distinctly from control samples (grey) along principal component 1 (PC1) at 4 h (A) and 24 h (B), indicating significant differences in their expression profiles. Each point represents a biological replicate.

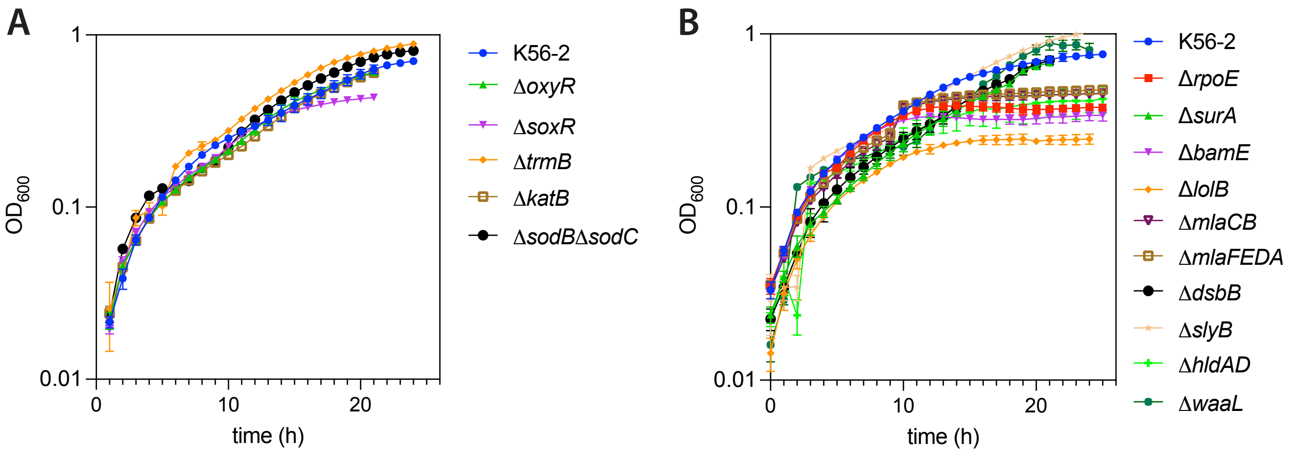

**Fig S2**. Growth kinetics of *B. cenocepacia* K56-2 and mutant strains in LB medium. **(A)** Growth curves of mutants lacking genes involved in the oxidative stress response. **(B)** Growth curves of mutants lacking genes associated with envelope stress response and cell envelope integrity.

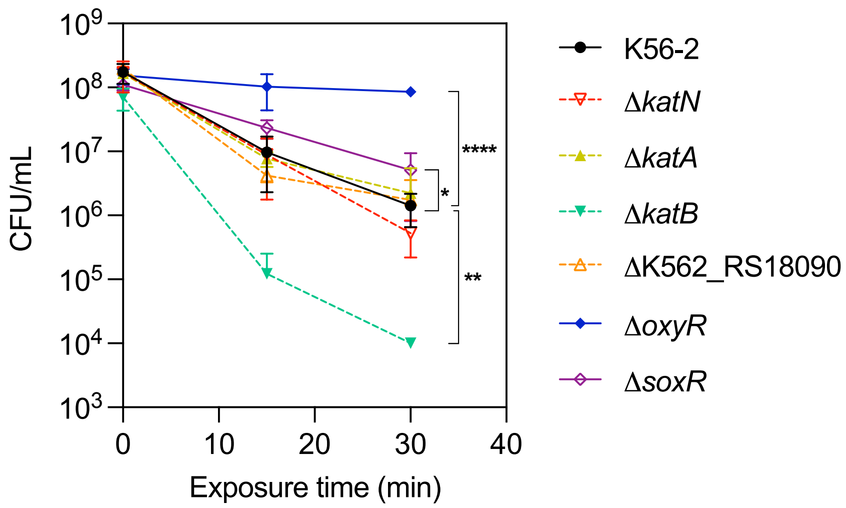

**Fig S3**. Survival curves of wild type strain K56-2 and mutants deficient in oxidative stress response genes following exposure to 4.26 mM $H_2O_2$ in LB medium. At selected time points (0, 15 and 30 min), bacteria were plated on LB and surviving bacteria were quantified by CFU counts. Data shown correspond to the average value from three independent biological replicates. Significant differences (* $P < 0.05$, ** $P < 0.01$, **** $P < 0.0001$) between mutants and K56-2 for each time point were determined using unpaired Student's t-test.

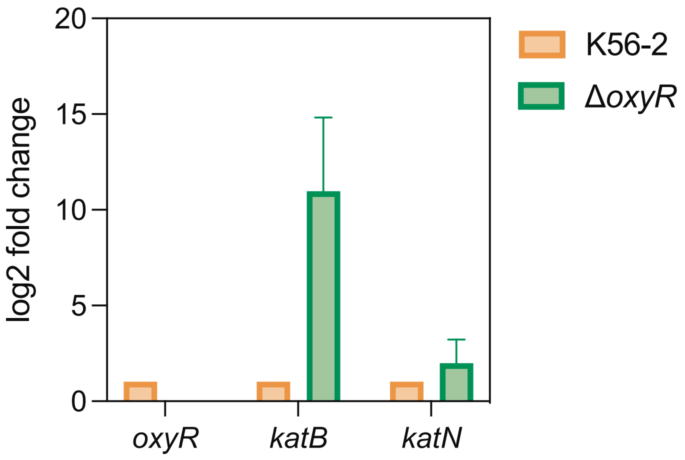

**Fig S4**. Expression levels of *oyxR*, *katB* and *katN* in the K56-2 wild type and Δ*oyxR* mutant strains. Transcript levels were quantified by RT-qPCR using RNA extracted from each strain growing in LB medium (log phase cultures, $OD_{600}$ = 0.3). Expression levels were normalized to the housekeeping gene *gyrA* and are shown relative to those in the K56-2 strain. No *oxyR* transcript was detected in the Δ*oxyR* strain, consistent with the deletion of *oxyR* gene in this mutant.
